# Petal photosynthesis and dynamic alterations of carbon metabolism orchestrate floral maturation in *Gardenia carinata*

**DOI:** 10.64898/2026.08.31.748199

**Authors:** Rohit Ghosh, Adinpunya Mitra

## Abstract

During maturation, flowers exhibit complex and finely-coordinated metabolic events facilitating various physiological changes that occur in a short time-span. The floral carbon metabolism underpins a metabolic frame, which provides energy and metabolites to drive the life-processes in flowers. However, studies on floral maturation in the past mostly focused on specialized metabolism or certain metabolite classes of central carbon metabolism. This study aims to provide a detailed insight into the dynamics of floral carbon metabolism and its role in flower maturation; it also addresses a long-standing question on the role of petal photosynthesis in floral maturation. In this report, time-course metabolomic studies were integrated with various physiological, enzymological, histochemical and ultrastructural findings to present a multi-layered investigation on the maturation physiology of *Gardenia carinata* flowers. This integrated approach revealed the characteristic metabolic features of different floral maturation stages, and also highlighted the metabolic rearrangements that allow each stage to establish their characteristic metabolic state. The photosynthetic bud stages showed metabolism inclined toward active growth and carbon reserve accumulation. A shift to energy-rich metabolism was observed as the flowers unfurled, probably to support energy-demanding events such as flower opening and changes in specialized metabolism. Our study further demonstrates the mechanisms underlining the transition from autotrophic to heterotrophic metabolism during maturation, which also reflects the plasticity in source-sink dynamics of flowers. We envisage that this study could provide a springboard for elucidating the roles of individual metabolites and enzymes in regulating floral maturation.

**Highlight:** Dynamic changes in floral carbohydrate metabolism enable the flowers to undergo a rapid physiological change in a short time-frame, where petal photosynthesis plays a pivotal role.

## Introduction

Flowers are ephemeral structures that arise as floral buds. During maturation, a well-orchestrated physiological and metabolic changes allow the buds to develop and unfurl into fully opened flowers (Borghi *et al*., 2022). The changes in central carbon metabolism that occur during flower opening enable them to display their reproductive potential through biosynthesis of different pollination cues such as scent volatiles and floral pigments. Besides, carbon metabolism in opened flowers also provide the energy and metabolites needed for nectar synthesis and secretion (Muhlemann *et al*., 2012; Dudareva *et al*., 2013; Ghissing *et al*., 2025). A gradual depletion of carbon reserve is observed following the opening of flowers, which culminates in metabolic and physiological rearrangements that end the floral lifespan (Ghosh *et al*., 2026). Understanding these dynamic changes in carbon metabolism thus becomes crucial to elucidate the physiology governing maturation and longevity of flowers.

Flowers are traditionally considered as non-photosynthetic organs, and our idea conventionally remain biased toward the assumption that they are completely heterotrophic and solely rely on the assimilates supplied from their mother plants. Nevertheless, petals constitute a non- reproductive whorl that produce cues for enticing pollinators and thus considered energetically and metabolically expensive to the mother plants (Stead *et al*., 2006). Study on *Nigella* spp. showed removal of the perianth resulted in increased biomass of the seeds, explaining the high energy demand of these whorls (Andersson, 2000; Andersson, 2005). However, petals of many flowers contain chlorophyll in the bud stages indicating their photosynthetic abilities (Pyke and Page, 1998; Ghosh *et al*., 2026). Presence of chlorophyll in the early stage of petal maturation and its subsequent degradation in the latter stages suggest petals as a site for mixotrophic metabolism, where early bud stages could be autotrophic and late maturation stages are heterotrophic (Paul *et al*., 2026). A study on *Nicotiana tabacum* flowers revealed the presence of Rubisco enzyme in the young petals, which supports active carbon assimilation. However, late maturation stages of *N. tabacum* petals showed reduced photosynthetic activities (Müller *et al*., 2010). Another study on *Gardenia jasminoides* revealed similar observations, where green buds showed the features of photosynthesis, which were absent in the latter stages (Ghosh *et al*., 2026). Though these studies mostly demonstrate that the petals possess features associated to photosynthesis, a detailed insight into the photosynthetic abilities of petals and how this impact floral maturation remained elusive. A comprehensive investigation integrating petal photosynthesis with physiological and metabolic changes that occur throughout floral maturation is therefore necessary to understand how petal photosynthesis contribute to this process and also to re-evaluate the existing prejudice regarding the heterotrophicity of flowers.

Though the view on trophicity of floral buds remains incompletely understood and advocates investigations, a few attempts have been made to understand central metabolism in flowers. Broadly, flowers are considered to act as sink that import sucrose as carbon source from mother plants for producing the energy and metabolites needed for running the physiological processes (Muhlemann *et al*., 2012). In flower maturation, the initial phase is represented by floral buds, which show active growth and gradual accumulation of storage reserve. During the growth phase, buds gradually increase in their dimensions and biomass (Dar *et al*., 2014; Önder *et al*., 2022; Jhanji *et al*., 2023). Such change is associated with the accumulation of storage compounds namely starch. Studies on *Rosa* spp. showed that petals gradually accumulate a substantial amount of starch during bud maturation (Kumar *et al*., 2008; Önder *et al*., 2023a). However, after anthesis, a decline in starch content was observed in many flowers (Yap *et al*., 2008; Mitra *et al*., 2025). The degradation of storage polysaccharides was shown to occur when buds transitioned into fully bloomed flowers (Ghosh *et al*., 2026). The released monosaccharides provide the energy and metabolites for biosynthesis and emission of floral scent, pigment accumulation and nectar secretion (Kutty *et al*., 2021; Ghissing *et al*. 2025, Paul *et al*., 2026). A few studies showed increased activities of sucrose metabolizing enzymes viz. sucrose synthase (SUS) and invertases (INVs) in petals upon flower opening. These ensure a supply of sucrose needed to supplement the existing storage reserve to fulfill metabolic demand of the petals (Koch, 2004; Stein and Granot, 2019; Borghi *et al*., 2022). As the flowers move towards senescence, major rearrangements in floral metabolism were observed (Kutty *et al*., 2021; Borghi *et al*. 2022). As floral senescence proceeds, the cell wall materials started to degrade and released monomeric building blocks such as glucose, possibly to fuel the terminal developmental process (Önder *et al*., 2023b). Interestingly, it was argued that the senescing flowers might rapidly remobilize the remaining carbon resources to actively growing young buds of the inflorescence, possibly for fulfilling their carbon demand (Bieleski, 1995; Sklensky and Davies, 2011; Borghi and Fernie, 2017). Collectively, these studies demonstrate that several metabolic rearrangements take place during floral maturation, and enables the flowers to undergo rapid physiological changes in a short time-frame. From the literature it was evident that existing studies mostly focused on selected aspects of floral metabolism and thus an integrated study demonstrating the dynamics in carbon metabolism is required to obtain a detailed insight into floral metabolism.

*Gardenia carinata*, a rubiaceous shrub, is popularly cultivated for their colour changing fragrant flowers. The flowers unfurl during evening and appear white. Gradually the flowers change their colour from white to saffron in a span of three to four days. In this report, we have presented a comprehensive account on the carbon metabolism of flowers in general, and of *G. carinata* in particular. To obtain a deep understanding of floral metabolism and to re-evaluate the trophic nature of flowers, we studied photosynthetic capacities and associated metabolic reprogramming of petals across the maturation stages. In addition, we combined semi-targeted metabolomic analyses with several physiological, enzymatic and microscopic studies to capture the dynamic changes in carbon metabolism associated with flower maturation. To elucidate the source-sink dynamics of petals, emphasis was given on sucrose metabolism, which also substantiates the idea on trophic status of flowers. In addition, through a photosynthesis perturbation experiment, we investigated a long-standing question on the role of early petal photosynthesis in floral maturation (Brazel and Ó Maoiléidigh, 2019). Broadly, our experiments provide insights on the metabolic mechanisms underlying the physiological changes associated with each floral maturation stage. The photosynthesis perturbation experiment further demonstrated that bud photosynthesis is essential for the accumulation of carbon reserve during bud maturation, and its disruption might cause reduced lifespan in *G. carinata* flowers. Together, our findings provide a holistic model that advances our existing understanding of metabolic basis of floral maturation. We envisage that our study will provide a springboard for elucidating the roles of specific metabolites, metabolic routes or enzymes that govern flower maturation and in turn its reproductive fitness.

## Material and Methods

### Plant material

*Gardenia carinata* Wall. ex Roxb. plants were grown in the experimental garden the department; identity of the species was confirmed by the Botanical Survey of India (CNH/Tech.II/2019/64 dated 7^th^ October 2020). The plants were grown under natural light conditions (*ca.* 42,000 Lux at 12:00 h). We studied flowers of six consecutive maturation stages which are as follows: early bud stage (S1: 6-7 days before anthesis), mid bud stage (S2: 3-4 days before anthesis), mature bud stage (S3: 1 day before anthesis), anthesis flower (S4: 0 h of flower opening), post-anthesis flower (S5: 1 day after flower opening) and senescent flower (S6: 3 days after flower opening).

### Biomass measurement

Fresh and dry petal mass was calculated as mentioned by Goswami and Mitra (2023). First, the fresh tissues were weighed and fresh mass was recorded. Then the tissues were lyophilized and weighed again to determine dry mass.

### Relative water content

Procedure mentioned by Önder et al. (2022) was followed for determining relative water content (RWC). Seventy petal discs (made from freshly plucked flowers using a cork borer) were weighed and were immersed in 25 mL deionized water for 4 h at 27 . Then the petal disks were blotted dry and weighed to obtain their turgid mass. The petal disks were then freeze-dried and their weight was recorded to get the dry mass. The following formula was used for calculating RWC:

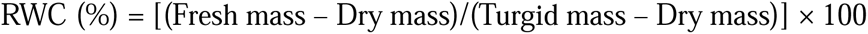

### Estimation of electrolyte leakage

Electrolyte leakage was estimated by following the method of Chen et al. (2008). Seventy discs of 50 mm diameter were made from petals using a cork borer. The petal discs were immediately immersed in 25 ml deionized water for 30 min, and initial conductivity (A1) of the solution was recorded using a conductivity meter (Hanna Instruments, USA). The solution containing petal discs was then heated to 100 for 30 min and final conductivity (A2) was recorded. Electrolyte leakage (%) was calculated by the formula: (A1/A2) × 100.

### Determination of cell integrity

Evans blues staining was used for determining the cell integrity following Delledonne et al. (2001) after minor modifications. The floral tissues were incubated in Evans Blue solution (0.2% w/v) for 3 h. The samples were washed thoroughly with deionized water followed by solubilization of Evans Blue molecules bound with dead cells in ethanolic solution (50% v/v) containing 1% (w/v) SDS. Absorbance of the obtained solutions were taken at 600 nm. Relative absorbances were plotted taking S1 stage as control.

### Measurement of Hill reaction activity

Protocol described by Tuohy and Choinski (1990) was followed for extraction of chloroplasts from the petal tissues, and 2, 6 dichlorophenolindophenol (DCPIP) method was followed for measuring Hill activity (Goswami and Mitra, 2023). The reaction mixture consisted of 300 µl 100 mM phosphate buffer (pH 7.0), 125 µl chloroplast extract and 75 µl of 3 mM DCPIP solution. The reaction mixture was incubated under light (*ca.* 3500 Lux) for 15 min and absorbance of the mixture was recorded at 600 nm in every 3 min interval. The blank was prepared by adding 125 µl extraction buffer in place of chloroplast extract.

### Measurement of chlorophyll fluorescence parameters

Chlorophyll fluorescence parameters were measured using a fluorometer (FluorPen FP110, Photon Systems Instruments, Czech Republic). The petals were dark adapted for 20 min with custom clips provided with the instrument before taking the measurements (Müller et al., 2010).

### Estimation of chlorophyll content

Protocol mentioned by Goswami and Mitra (2023) was followed for extraction of chlorophyll. At first, fresh petal tissues were homogenized in 80% (v/v) acetone and the homogenate was centrifuged at 5000 *g* for 7 min. Chlorophyll quantity was estimated following the method described by Lichtenthaler (1987).

### Histochemical analysis of floral petals

Histochemical analysis was performed to study the localization of metabolite accumulation patterns inside the petal tissues across the maturation stages. The petal tissues were cut into 2×2 mm pieces and cryoprotected using aqueous solution of glycerol (10%, v/v) at 4 . The cryoprotected floral tissues were then sectioned at a thickness of 20 µm using a semi-automatic cryostat microtome MCM-MT (Medimeas Instruments, Haryana, India) (Paul and Mitra, 2024). To detect the presence of chlorophyll, the sections were excited with UV-light (365 nm) to capture the resulting fluorescence. The sections were stained with Lugol’s iodine and periodic acid-Schiff (PAS) reagent for detection of starch and polysaccharides, respectively (Paul and Mitra, 2024). The sections were observed in a Leica^TM^ DM2500LED upright microscope and photographed with Leica^TM^ DFC7000T CCD camera using Leica Application Suite X platform (Leica, Germany).

Histolocalization of glucose was done in fresh sections following the protocol of Martinelli (2008) with slight modification. The sections were incubated for 30 min in 80 mM Tris-HCl buffer (pH 7.8) containing 10 mM EDTA, 20 mM MgCl_2_, 0.8 mM NAD, 0.6 mM ATP, 10 U ml^-1^ glucose-6-phosphate dehydrogenase, 20 U ml^-1^ hexokinase and 0.03% iodonitrotetrazolium chloride (INT).

For histolocalization of enzymes, the petal samples were cut into 2×2 mm pieces and fixed in 50 mM phosphate buffer saline containing 2 mM DTT, 2% (w/v) paraformaldehyde and 2% (w/v) polyvinylpyrrolidone (PVP) at 4 for 1 h (Sergeeva and Vreugdenhil, 2002; Baud and Graham 2006). After fixation, the samples were washed five times and put in reaction buffer. For histochemical assays performed in petal transverse sections, the samples were first cryoprotected as mentioned previously and washed with water (Sergeeva and Vreugdenhil, 2002). Cell wall invertase was histochemically assayed following the protocol mentioned by Sergeeva and Vreugdenhil (2002) and Paul and Mitra (2024). The obtained petal sections were re-fixed using the same fixative buffer mentioned earlier. The cross-sections were incubated in a 0.038 M phosphate buffer (pH 6.0) containing 25 U glucose oxidase, 1% (w/v) sucrose, 0.024% (w/v) NBT and 0.014% phenazine methosulfate for 8 h and observed using microscope. For determining enzyme activities of ADP-glucose pyrophosphorylase (AGPase) and UDP-glucose pyrophosphorylase (UGPase) in petal tissues though histochemical techniques, protocol mentioned by Jammer *et al*. (2015) were followed and coupled with NBT for localization of enzyme activity (Jammer *et al*., 2020). The samples were incubated with staining buffer containing 5 mM MgCl_2_, 0.44 mM EDTA, 0.1% BSA, 1.5 mM PPi, 2 mM 3-PG, 1 mM NADP, 1.3 U of G6PDH, 0.45 U of PGM, 0.8 mM NBT and 2 mM ADPGlc (for AGPase) or UDPGlc (for UGPase) in 100 mM Tris buffer (pH 8.0). The control reactions were devoid of ADPGlc and UDPGlc. Protocols mentioned by Sergeeva and Vreugdenhil (2002) and Baud and Graham (2006) were followed for detection of phosphoglucomutase (PGM), phosphoglucose isomerase (PGI) and hexokinase (HK) activities. For detection of PGM activity the fixed plant samples were incubated for 4 h in 0.042 M HEPES buffer (pH 7.4) containing 0.84 mM EGTA, 0.084 mM EDTA, 0.08% BSA, 4.2 mM MgCl_2_, 1.5 mM NAD, 0.4 mM NBT, 1.2 U ml^-1^ G6PDH and 4.5 mM glucose-1-phosphate (G1P). For detection of PGI activity, in place of G1P, 4.5 mM fructose-6-phosphate was added. The HK activity was determined by incubating the fixed petal samples for 3 h in 0.05 M Bis-Tris buffer (pH 8.0) containing 2.5 mM ATP, 0.025% (w/v) BSA, 6.25 mM MgCl_2_, 1.2 U ml^-1^ G6PDH, 1.5 mM NAD, 0.4 mM NBT, 0.25 mM EDTA, 0.25 mM EGTA and 0.5 mM glucose. The petal tissues were analyzed using Fiji (based on ImageJ 1.54p) for quantification purposes.

### Semi-targeted analysis of petal metabolites

Lyophilized petal tissues were used for extraction and subsequent analysis of petal metabolites. The lyophilized tissues were powdered using liquid N_2_ in a mortar and pestle. The powdered samples (5 mg) were extracted in methanol (1 ml containing 100 µg ribitol) at 70 for 15 min. A small part (100 µl) was aliquoted in a microcentrifuge tube and vacuum dried before derivatization. The dried samples were methoxyaminated using methoxyamine-HCl (30 mg ml^-1^ pyridine) at 37 for 120 min followed by silylation using N, O- bis(trimethylsilyl)trifluoroacetamide at 70 for 30 min (Kotamreddy *et al*. 2020). For analysis, 1 µl of derivatized samples were injected in GC-MS. GC-MS parameters are detailed in supplementary method S1.

### Analysis of apoplastic metabolites

Collection of petal apoplastic metabolites were done following the protocol of O’Leary *et al*. (2016) using a vacuum flask. Freshly collected petals were rinsed in deionized water and blot-dried to clean the petals. Petals were then kept in a modified conical flask containing deionized water and vacuum was applied. The flask was slowly agitated for 5 min and slowly the vacuum was released. The vacuum cycle was repeated until the petal tissues were completely infiltrated with water. The petals were then removed from the flask, gently blotted and then rolled inside a parafilm strip. The rolled petals were then placed inside modified pipette tips and the tips were placed inside centrifuge tubes. The samples were then centrifuged at 600 *g* for 10 min. Dilution of collected apoplastic fluids were calculated using indigo carmine. Apoplastic metabolites were analyzed in GC-MS after derivatization following the protocol mentioned by Kotamreddy *et al*. (2020) as described earlier. Here, apoplastic fluid was added in methanol instead powdered plant sample.

### Estimation of cellulose quantity

Protocol mentioned by Yap *et al*. (2008) was used for estimation of cellulose. First, fresh floral tissued were homogenized using chilled 95% (v/v) ethanol and kept overnight at -20 . On the next day, the tubes were centrifuged at 8500 *g* for 12 min at 4 . Tris buffer (0.5 M, pH 7.5) containing phenol was then added in the pellets, vortexed and incubated for 45 min at room temperature. The samples were again centrifuged and 80% (v/v) ethanol was added to the pellets and kept at -20 for 3 h. The samples were then centrifuged and washed with 80% (v/v) ethanol, 80% (v/v) acetone and 1:1 mixture of methanol-chloroform. The obtained residue was filtered and washed three times with acetone. Finally, the obtained residue was dried and 10 mg of the sample was hydrolyzed using 3 ml of 10:1 mixture of acetic acid and nitric acid for 30 min at 100 . Then the samples were re-centrifuged, washed with water and finally, incubated for 1h with 10 ml of 67% H_2_SO_4_. Cellulose quantification from the hydrolyzed samples were performed using anthrone method. Cellulose was used for preparation of the calibration curve.

### Estimation of starch quantity

Protocol mentioned by Bahdanovich *et al*. (2022) was used for determination of starch quantity in the petal tissues. At first, powdered plant samples were heated at 105 for 24 h and 1 ml of 1 M NaOH and 100 µl of 95% (v/v) ethanol was added before incubation at 4 for 24 h. Then, the final volume of the mixture was maintained till 10 ml and again kept at 4 for 16 h. Finally, pH of the samples were maintained at 6.0 with HCl. Starch quantities of the samples were determined by adding 0.2% (w/v) KI/I_2_ solution with the extracts. Absorbance caused by amylose-iodine conjugation were taken at 590 nm. Starch was used for preparation of the calibration curve.

### Estimation of fructan quantity

For determination of fructan quantities, protocol mentioned by Yap *et al*. (2008) was followed. At first, 200 mg of plant samples were boiled for 90 min to remove soluble sugars. Then, fructans were extracted from the petal tissues in 10 mM acetate buffer (pH 4.5) by incubating the solutions for 24 h at 30 . Finally, the samples were centrifuged at 7500 *g* for 15 min and supernatants were collected. Phenol-sulphuric acid method was used for spectrophotometric detection of fructans from the obtained supernatants.

### Determining *in vitro* activities of enzymes involved in cell wall breakdown

Crude protein extraction from the floral tissues were performed following the protocol mentioned by Roy Choudhury *et al*. (2009). Freshly collected 200 mg tissue was homogenized in a chilled 25 mM HEPES buffer (pH 7.5) containing 5 mM β-mercaptoethanol, 5 mM MgSO_4_, 2 mM DTT and 15 mM KCl. The homogenized floral tissues were centrifuged at 12000 *g* at 4 for 15 min. The collected supernatants were used as the source of crude proteins. Cellulase and xylanase enzyme assays were performed following the protocols mentioned by Srivastava and Dwivedi (2000). The reaction mixture contained 100 mM acetate buffer (pH 5.0), crude extract and 1% (w/v) carboxymethylcellulose (for cellulase) or 0.1% beechwood xylan (for xylanase) and incubated for 4 h for cellulase and 2 h for xylanase. Glucanase activity was determined by following the protocol mentioned by Mukherjee *et al*. (2023), where the reaction mixture contained 50 mM acetate buffer (pH 4.8), crude extract and 2% (w/v) laminarin and was incubated for 1 h. Product formed in these enzyme assays were determined by DNS method and enzyme activities were expressed with respect to the formed product (Srivastava and Dwivedi, 2000).

### Determining *in vitro* activities of enzymes involved in starch breakdown

For determining starch degrading α-amylase activities, protocol mentioned by Ghosh *et al*. (2026) was followed. Fresh petal tissues were ground using liquid nitrogen and immediately transferred to 0.1 M acetate buffer (pH 5.5) containing 3 mM calcium chloride and 1 mM β- mercaptoethanol in microcentrifuge tubes and vortexed for 10 s. Then the tubes were cold centrifuged for 20 min at 12000 *g*. Supernatants were collected for enzyme assays. The reaction mix contained 40 µl enzyme extract, 3 mM calcium chloride, 1 mM β-mercaptoethanol, 8% (w/v) soluble starch in 0.2 M acetate buffer in a total reaction volume of 200 µl. The reaction tubes were incubated for 30 min at 37 . Formed product quantities were determined using DNS method for calculating enzyme activity.

For determining *in vitro* β-amylase activities, crude protein extraction and assay were conducted following Guleria and Kumar (2023) with minor modifications. The petal tissues were ground using liquid nitrogen and transferred immediately to 0.1 M Tris-HCl buffer (pH 6.5) containing 8 mM MgCl_2_, 0.1 mM phenylmethylsulfonyl fluoride (PMSF), 1 mM DTT and 2 mM EDTA in microcentrifuge tubes and vortex for 10 s. The slurries were then cold centrifuged for 15 min at 10000 *g*. The collected supernatants were used for enzyme assays. Crude protein extracts were mixed and incubated with reaction mixture containing 0.78 mM EDTA and 5% (w/v) soluble starch in citrate buffer (50 mM, pH 3.6) for 1 h at 24 . Formed product concentrations were determined using DNS method for calculating enzyme activities.

### Determining *in vitro* activities of sucrose metabolizing enzymes

For determining the sucrose phosphate synthase (SPS) and sucrose synthase (SUS) enzyme activities, protocols mentioned by Paul *et al*. (2026) were followed with slight modification following Jammer *et al*. (2020). Fresh petal tissues were extracted in chilled 50 mM HEPES-KOH buffer (pH 7.5) containing 2% (w/v) PEG 6000, 5 mM MgCl_2_, 1% (w/v) PVP, 2mM EGTA, 1mM β-mercaptoethanol and 2.5 mM DTT. The extracts were centrifuged in refrigerated condition at 17500 *g* for 15 min. The collected supernatants were used as source of enzymes. The reaction mixture contained 50 mM MgCl_2_ in 200 mM HEPES-KOH buffer (pH 7.5). With the reaction mixture, 25 mM UDP-glucose and 25 mM fructose-6-phosphate (for SPS) or 25 mM UDP-glucose and 25 mM fructose (for SUS) were added as enzyme substrates. Protein extracts were added to start the reactions. Invertase (INV) assays were performed following the protocol mentioned by Jammer at al. (2015). Reaction mixture contained crude protein extract, 25 mM sucrose solution in buffers of pH 6.8 (114 mM citric acid/772 mM Na_2_HPO_4_) or pH 4.5 (273 mM citric acid/454 mM Na_2_HPO_4_) for cytosolic invertase (cINV) or vacuolar invertase (vINV), respectively. Formed product concentrations were determined for calculating enzyme activities.

### Photosynthesis perturbation in petals

The S1 stage petals were covered in sleeves made with black cloth which ensures almost complete blocking of sunlight. The buds were grown till S3 stage and the experiments were conducted. Normally growing buds were taken as control in the experiments.

## Statistical analysis

At least three imprecise biological replications were taken for all experiments. The plots show means of obtained values and the error bars represent standard deviations. For calculating statistical differences, t-test and one way ANOVA with Tukey’s HSD were performed. Statistical significance of values was considered at *P* ≤ 0.05. MetaboAnalyst 6.0 platform was used for statistical analysis of metabolomics data.

## Results

### Study of phenotypic changes in different maturation stages of *G. carinata* flowers

This study focused on comprehensive understanding of changes in central metabolites that occur in *G. carinata* flowers during maturation through multifaceted approaches. During flower maturation, the buds mature and unfurl into white flowers, which subsequently change their colour to saffron as they approach senescence (Fig. 1A). A gradual increase in fresh and dry mass of petals was observed as the flowers progress through S1 to S4; a subsequent decline of biomass was noticed till S6 (1B, C). Highest relative water content was observed in S4 and S5, and a significant decline was recorded from senescent flowers thereafter (1D). Loss in tissue integrity and cell death in petals were assessed by measuring electrolyte leakage and Evans Blue staining, respectively. A gradual increase in electrolyte leakage was observed after anthesis, with the S6 petals showing highest electrolyte leakage (1E). Likewise, highest Evans Blue accumulation was observed in S6 stage, indicating cell death in the senescent petals (1F).

**Fig. 1.**
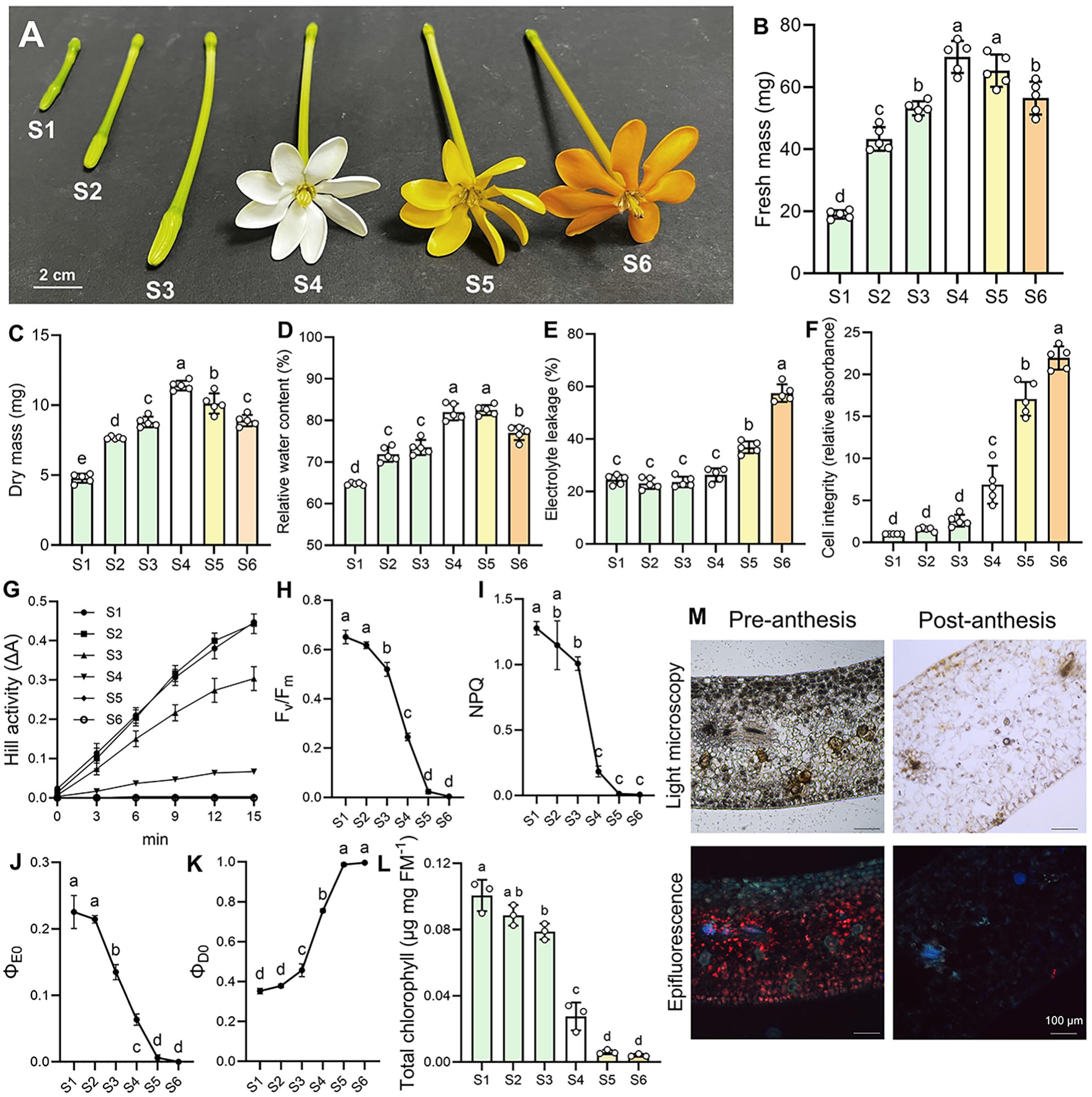
Phenotypic and photosynthetic changes in *G. carinata* flowers during maturation. Different stages of floral maturation stages studied here is shown in the composite image (A). Fresh and dry masses of *G. carinata* flower petals (B, C). Changes in relative water content during floral maturation (D). Changes in electrolyte leakage and cell integrity (E, F). Estimation of Hill activity (G), maximum quantum yield of PSII (F_v_/F_m_) (H), non-photochemical quenching (I), quantum yield of electron transport (Φ_E0_) (J) and quantum yield of energy dissipation (Φ_D0_) (K). Quantification of total chlorophyll (L). UV-excitation on cryo-sections showing presence of chlorophyll (red fluorescence) towards the abaxial surface of pre-anthesis petals; no autofluorescence (red) is found in post-anthesis abaxial petal surface indicating absence of chlorophyll (M). n ≥ 3 in all experiments. The bar charts represent mean of biological replicates ± SD and the dots show the data points. Statistical differences were determined by one way ANOVA and are indicated by different letters (*P* ≤ 0.05). Scale bars in the microscopic images represent 100 µm.

### Photosynthetic traits in *G. carinata* flowers

Leaves are the main photosynthetic organs of plants. However, other plant parts, e.g., stem, flowers and fruits can also synthesize a substantial quantity of photoassimilates. In such context, we have quantified various photosynthetic characteristics from *G. carinata* flowers. The highest Hill activity was recorded in S1 and S2 petals, followed by a significant reduction in S3 stage. The lowest Hill activities were recorded from anthesis and post-anthesis petals (S4 to S6) (Fig. 1G). In conjunction with Hill activity measurements, highest maximum quantum yield of PSII (*F*_v_/*F*_m_) was recorded in S1 and S2 stages, followed by a decline in S3 stage (Fig. 1H). Similarly, non-photochemical quenching (NPQ) showed significant reduction after the flowers unfurl (Fig. 1I). Quantum yield for electron transport (Φ_E0_) measurement also showed a comparable trend to that of *F*_v_/*F*_m_ and NPQ (Fig. 1J). However, quantum yield of energy dissipation (Φ_D0_) showed a opposite trend (Fig. 1K). We also found that the gas exchange rates in the petals of bud stages were significantly higher as compared to the other stages of floral maturation (Fig. S1). SEM micrographs of petal surface showed partial blockage of stomatal apertures with wax-like crystals in S4 and S6 stages (Fig. S2). These observations explain the occurrence of reduced gas exchange after flower unfurling. Spectrophotometric estimation of total chlorophyll showed highest chlorophyll content in S1 and S2 stages, and a gradual decline was observed in the latter stages of maturation (Fig 1L). Epifluorescence images also showed the presence of high abundance of chlorophylls towards the petal abaxial surface in pre-anthesis bud stages (Fig. 1M). TEM micrographs also showed the presence of chloroplasts in the petals of bud stage (Fig. S3).

### Changes in primary metabolism during floral maturation

To study the changes in primary metabolites in petal tissues, GC-MS based semi-targeted analyses were conducted across the floral maturation stages. The variations in metabolite concentrations are presented in a heatmap (Fig. 2A). Metabolite profiles of each maturation stages were hierarchically clustered to compare the stages, which grouped the floral maturation stages into two superclusters. The first supercluster nested the bud stages (S1 to S3) and the anthesis stage (S4). The bud stages were grouped into a common cluster, suggesting a similar metabolite composition of these maturation stages. In contrast, the anthesis flowers (S4) were present in a separate group of the supercluster, which indicates the occurrence of major reprogramming in metabolism as the flowers open. The flowers of S5 and S6 stages were present in the second supercluster. Hence, we can broadly assort the six maturation stages into three major maturation phases viz. pre-anthesis (S1 to S3), anthesis (S4) and post-anthesis (S5 and S6) (Fig. S4).

**Fig. 2.**
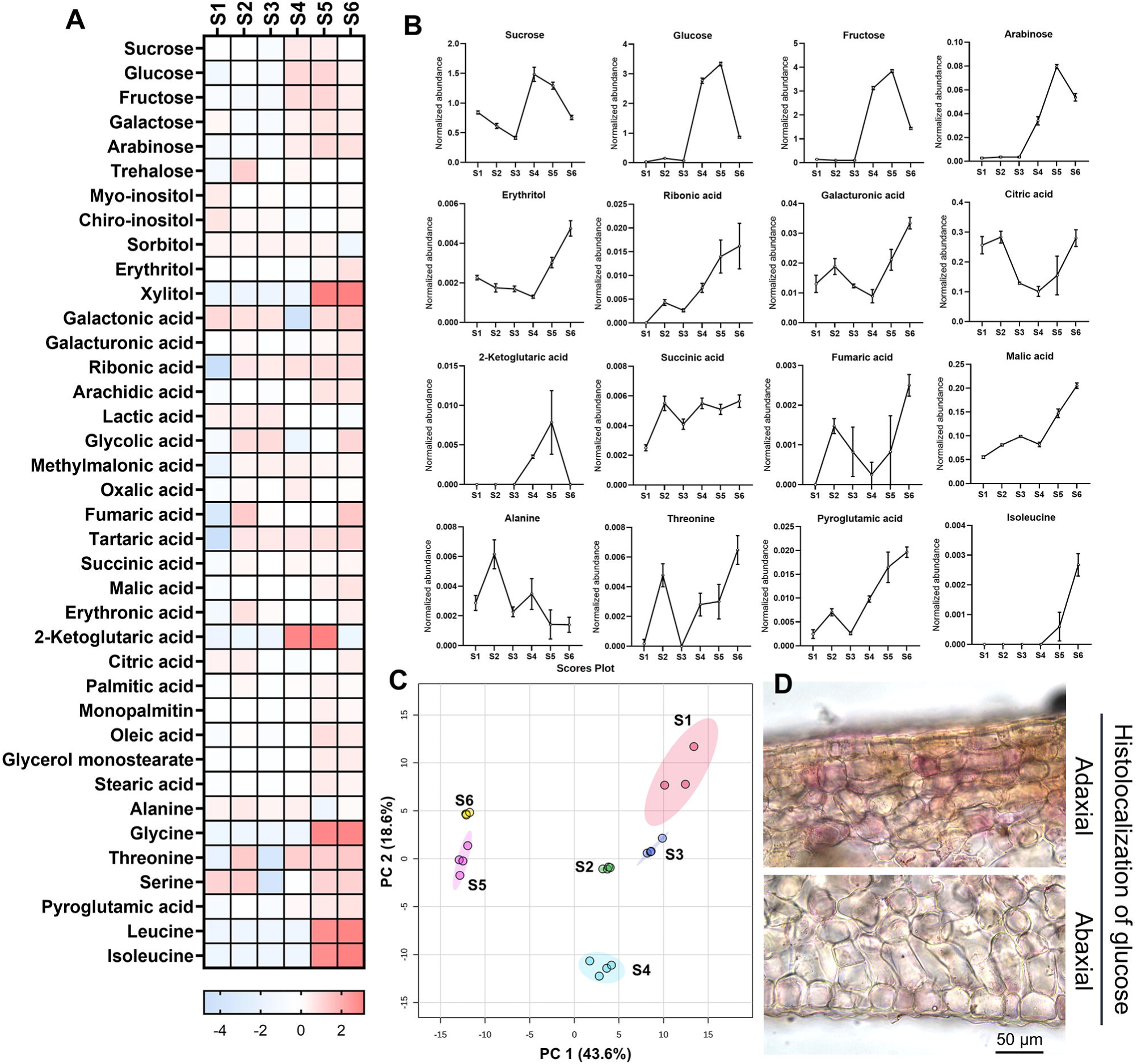
Variations in compositional diversity of central metabolites in *G. carinata* petals during maturation. Heatmap showing compositional variation of different metabolites across the floral maturation stages (A). Normalized abundance of metabolites was log_10_-transformed followed by pareto-scaling before heatmap construction. Normalization with internal standard was performed for calculating abundance. Abundance of various metabolites across the floral maturation stages (B). PCA scores plot representing 62.2% of total variation, depicting clear separation of the maturation stages in the scores plot (C). Histochemical analysis shows higher abundance of glucose towards the adaxial cell layers of petal, as observed by reddish-purple precipitation of reduced INT (D). n ≥ 3 in all experiments. Line graphs showing mean abundance of metabolites ± SD.

An increase in titer of metabolic sugars viz. sucrose, glucose and fructose were observed after the flowers unfurled (Fig. 1A, B). Highest quantities of glucose and fructose were recorded from S5 stage, characterized by pigments accumulation and highest volatiles emission (pigment and volatile data not shown). Trehalose concentration was highest in S2 bud followed by a sharp decline as the buds moved to S3 stage. An increase in trehalose concentration was observed in S4 flowers thereafter (Fig. 1A). Variable trehalose concentrations in different maturation stages are crucial to regulate starch and hexose metabolism. Sugar alcohols such as xylitol and erythritol showed highest concentrations in S5 and S6 stages, and sugar phosphates such as G1P and G6P concentration were found to be elevated in the growing bud stages (S1 to S3) (Fig. 2A; Fig. S5).

Assessment of petal metabolite profiles across the stages also revealed the variable content of different TCA cycle intermediates across the maturation stages. Citric acid and fumaric acid showed an initial increase in early bud stages and thereafter a decline was observed. Elevation in their concentrations were again observed in the senescent flowers (S6). 2-Ketoglutaric acid, succinic acid and malic acid showed highest concentration in S4 to S6 stages. Although the different TCA intermediates showed their distinct accumulation pattens, a common trend was their higher accumulation after flower opening (Fig. 2A, B).

We found a complex pattern of amino acid accumulation across the *G. carinata* floral maturation stages. Amino acid and their derivatives such as alanine, glycine, serine and pyroglutamic acid showed an initial increase, particularly in the actively growing S2 stage. Increased concentration of amino acids in the bud stage probably indicates active growth of the floral tissues. Later, in the post-anthesis S5 and S6 stages an increased concentrations of all amino acids were recorded. This observation suggests that degradation of protein probably causes release of amino acids during the terminal maturation stages (Fig. 2A). Highest abundance of fatty acids and their derivatives were recorded from post-anthesis S5 flowers (Fig. 2A). PCA analysis showed distinct separation of the floral maturation stages in the plot. The PCA plot could account for 62.2% of the total variation (Fig. 2C). Histochemical detection of glucose in cryo-sectioned petals of *G. carinata* flowers showed higher accumulation of glucose in the petal adaxial surface compared to the abaxial cell layers. This indicates the adaxial cell layers probably have higher metabolic activity to support the metabolite and energy demand for scent volatiles biosynthesis and release (Fig. 2D).

*In vitro* activities of glucose-6-phosphate dehydrogenase (G6PDH) showed significant increase upon flower opening (Fig. S6). Similarly, TCA cycle enzymes such as isocitrate dehydrogenase (ICDH) and succinate dehydrogenase (SDH) activity gradually increased after anthesis (S4 to S6) (Fig. S7, 8). However, glyceraldehyde-3-phosphate dehydrogenase (GAPDH) activity got significantly reduced after flower opening (Fig. S7).

### Metabolomic changes in apoplast

Apoplast is the extra-protoplasmic matrix of a cell including cell wall. The apoplastic space contains various metabolites and enzymes that modulate several physiological processes of plant organs. In order to study the apoplastic metabolites from petal tissues, three distinct physiological phases of floral maturation, i.e., pre-anthesis, anthesis, and post-anthesis stages into three distinct phases were selected, based on clustering of petal metabolomic data (Fig. S4). The collected apoplastic metabolites were derivatized, and their calculated abundance was represented in a heatmap. From the petal apoplast, we identified compounds belonging to different chemical classes including sugars, sugar alcohols, organic acids, fatty acids and amino acids. Metabolic sugars such as glucose, fructose, and sucrose showed highest abundance during anthesis, which indicate the relatively high metabolic activity in this scent-emitting stage (emission data not shown). We observed distinct trends for the different metabolites of TCA cycle. Citric and malic acids showed a progressive increase in their contents as the flowers mature, while fumaric acid showed an opposite trend. An initial increase of succinic acid concentration was noted during anthesis, which subsequently declined in the post-anthesis phase. These asynchronous concentrations of TCA cycle intermediates probably indicate the presence of different non-cyclic flux modes during floral maturation. Several amino acids such as glycine, alanine, isoleucine, etc. showed increased concentrations during the floral maturation (Fig. 3A, B).

**Fig. 3.**
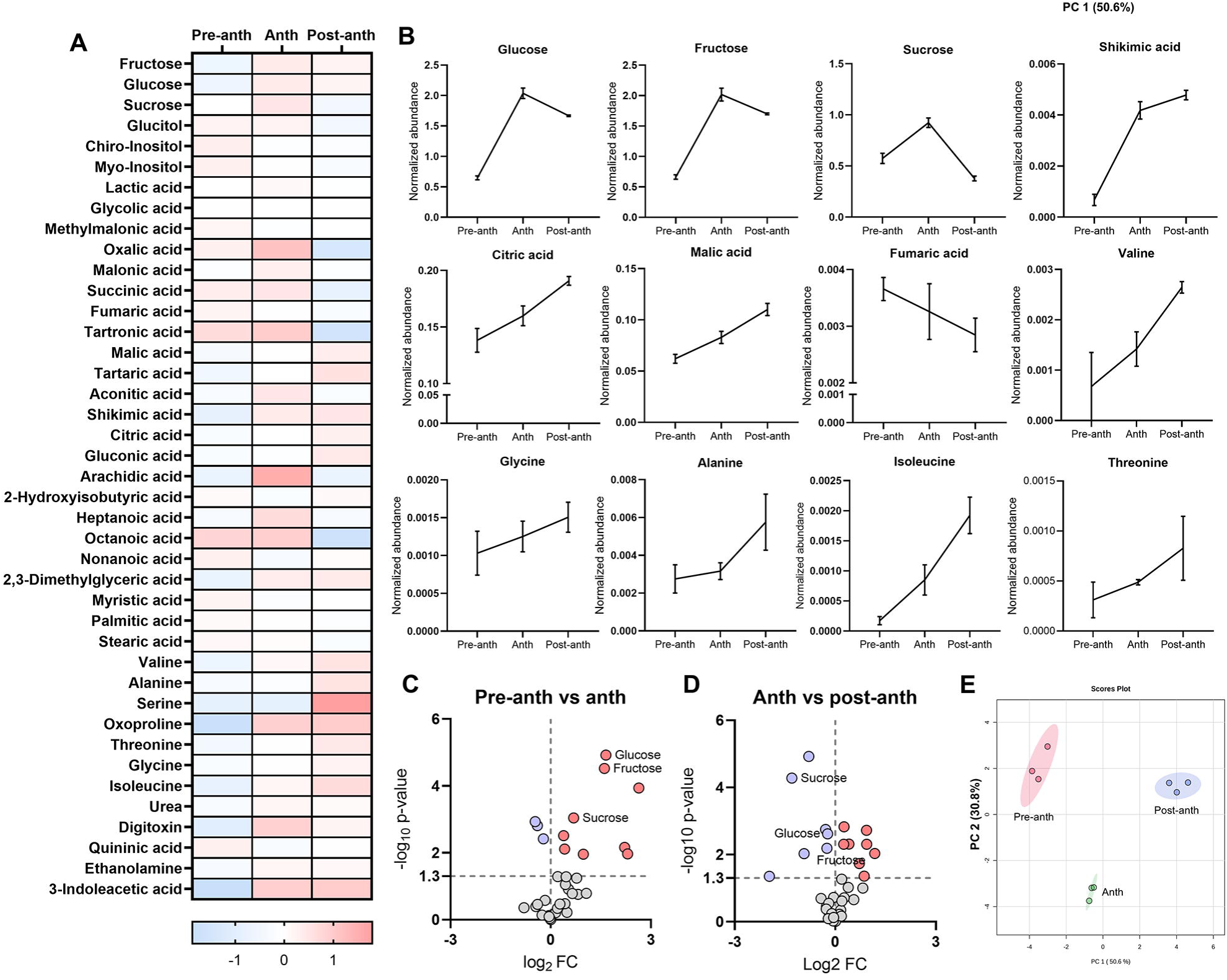
Variations in compositional diversity of apoplastic metabolites in *G. carinata* petals during maturation. Heatmap showing composition of apoplastic metabolites across the major floral maturation stages (A). Normalized abundance of metabolites was log_10_-transformed followed by pareto-scaling before heatmap construction. Normalization with internal standard was performed for calculating abundance. Variation of a few apoplastic sugars, organic acids, amino acids abundance across the maturation phases (B). Pairwise comparisons of metabolites are represented in volcano plots. Volcano plots indicate an upregulation in metabolic sugar contents (glucose, fructose and sucrose) during anthesis and eventual downregulation of the same in the post-anthesis petals (C, D). PCA scores plot showed a clear separation of the maturation phases along the first two components, indicating significant difference in metabolite composition (E). n ≥ 3 in all experiments. The bar charts represent mean of biological replicates ± SD.

Pairwise comparisons of the metabolomic data across the floral maturation phases are represented in two separate volcano plots (Fig. 3C, D). A significant upregulation of metabolic sugar content was observed when pre-anthesis buds transitioned to anthesis flowers, and subsequently a downregulation was observed as the anthesis flowers moved to the terminal maturation phase. PCA analysis showed distinct segregation among the clusters, indicating compositional uniqueness of the maturation phases (Fig. 3E).

### Studying the changes in cellulose, starch and fructan quantities

Cellulose, a complex carbohydrate made of glucose, is the most abundant polymer in the cell wall. We observed a gradual increase in cellulose content from the S1 to S4 stages. Subsequently, a decrease was observed as the post anthesis flowers move towards senescence (Fig. 4A). In TEM analysis we observed buckled and disintegrating cell wall in the senescing tissues. The depolymerization of wall components was observed progressing from the periphery towards the center (Fig. S3).

**Fig. 4.**
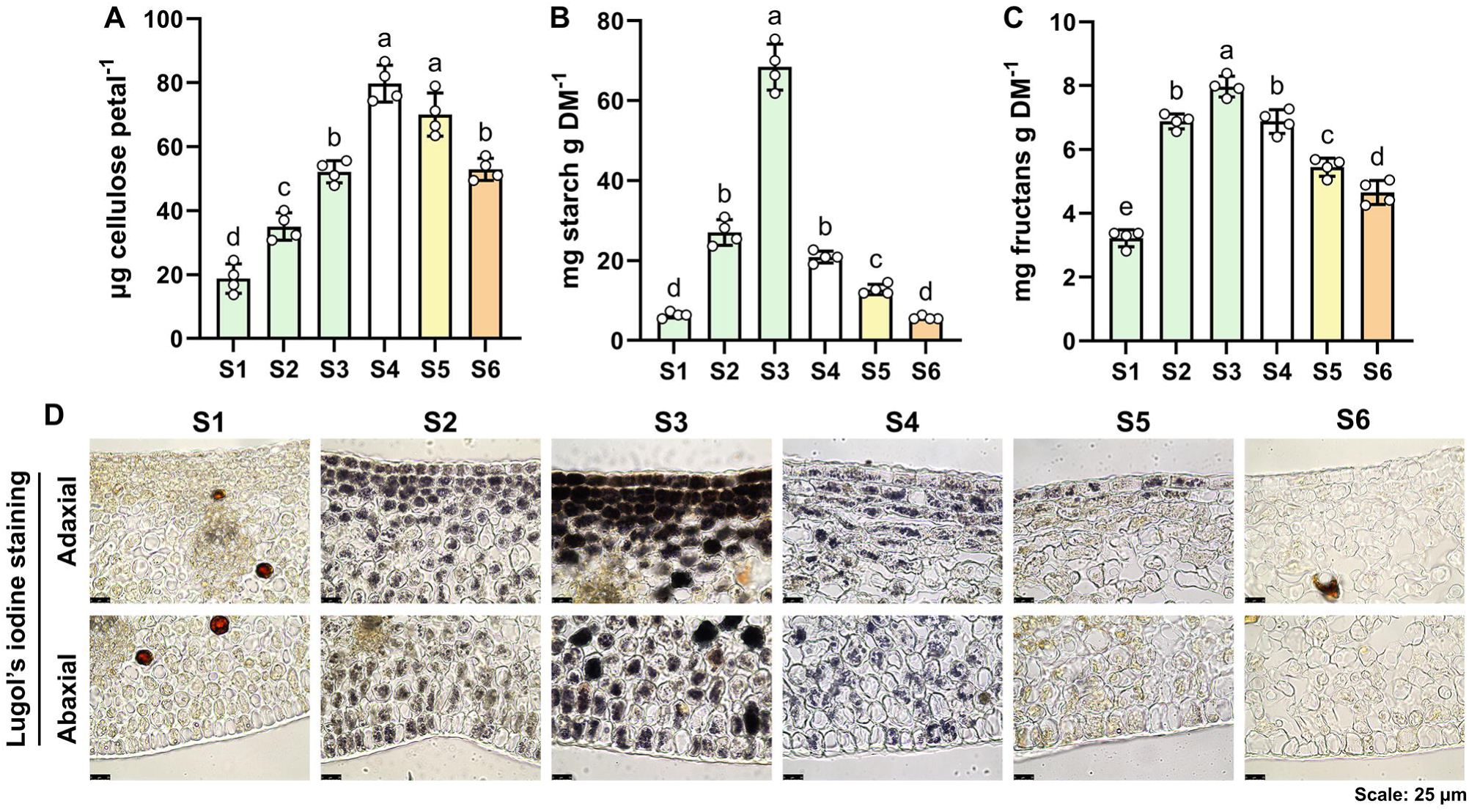
Estimation of cellulose, starch and fructan contents across different maturation stages of *G. carinata* flowers. Quantification of cellulose, starch and fructan contents in *G. carinata* flowers (A-C). The cryo-sections showing starch distribution patterns in petal tissues across different maturation stages (D). The sections were stained with Lugol’s iodine to detect starch granules, which appear dark brown in the micrographs. n ≥ 3 in all experiments. The bar charts represent mean of biological replicates ± SD and the dots show the data points. Statistical differences were determined by one way ANOVA and are indicated by different letters (*P* ≤ 0.05).

Starch and fructans function as primary reserve carbohydrates in plants. Quantities of the both increased as buds moved from S1 to S3 stage. A gradual decline in their quantities was observed as the flowers opened and transitioned towards senescence (Fig. 4B, C). Histochemical and TEM observations revealed a similar trend as found earlier (Fig 4D; Fig S4). In addition, we observed a higher starch accumulation in the adaxial cell layers of petal sections than the abaxial side at the S3 stage (Fig. 4D). A similar trend was found with PAS staining, which showed histiolocalization of carbohydrates within the petal tissues (Fig. S9).

### Activities of cell wall and starch metabolizing enzymes

Using histochemical techniques, activities of common enzymes involved in the both cellulose and starch biosynthetic routes such as HK, PGI, and PGM were detected. The enzymes showed significantly higher activities in the photosynthetic bud stages (S1 to S3) compared to the other maturation stages (S4 to S6). We also detected a higher activity of UGPase, an enzyme of the cellulose biosynthetic route in S1 to S3 stages, indicating active cell wall synthesis and growth in floral buds. However, AGPase - a key enzyme of starch biosynthetic pathway showed higher activities in S2 and S3 petals, and thereafter a significant decline was detected as the flowers moved through maturation (Fig. 5B, C). Similarly, starch synthase activity was higher in green bud stages, and declined thereafter (Fig. S6).

**Fig. 5.**
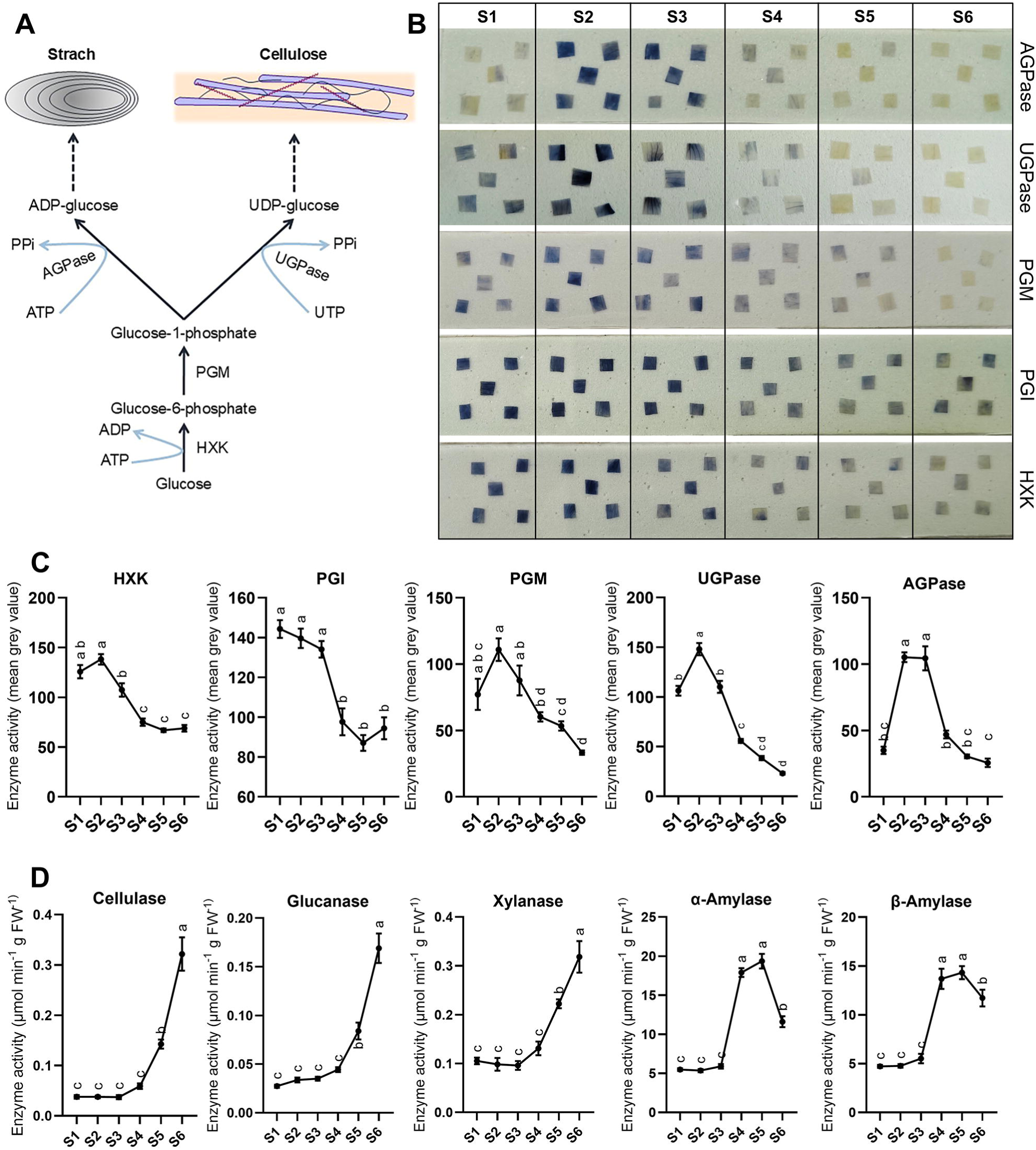
Estimation of enzyme activities associated with cell wall (cellulose) and starch metabolism. (A) A schematic representation of cellulose and starch biosynthesis pathway. (B) Images showing staining intensities in petal tissues after histoenzymological tests for different enzymes of starch and cellulose metabolism. (C) Line graphs showing activities of cell wall and starch biosynthesizing enzymes across different stages of floral maturation. (D) Line graphs showing activities of cell wall and starch degrading enzymes across different stages of floral maturation. n ≥ 3 in all experiments. The values corresponding to the maturation stages in the graphs represent the mean of biological replicates ± SD. Statistical differences were determined by one way ANOVA and are indicated by different letters (*P* ≤ 0.05). Abbreviations: HXK, hexokinase; PGI, phosphoglucose isomerase; PGM, phosphoglucomutase; UGPase, UDP-glucose pyrophosphorylase; AGPase, ADP-glucose pyrophosphorylase.

Floral maturation in latter stages, especially senescence, is characterized by disintegrating and thinning of cell wall. In this study, we assessed the activities of wall-degrading enzymes viz. cellulase, glucanase and xylanase, which respectively break down cellulose, glucans and xylans of the cell wall into simpler sugars. We found gradual increase in the activities of above enzymes after anthesis, which peaked in the senescent flowers (S6). Contrary to the activity trend shown by wall degrading enzymes, starch breakdown enzymes such as α and β-amylase showed elevated activities in S4 and S5 stages compared to the green stages, and afterward a decline was observed in S6 petals (Fig. 5D).

### Activities of sucrose-metabolizing enzymes

*In vitro* activities of sucrose-metabolizing enzymes including SPS, SUS and INVs were assessed across the stages of floral maturation. The activities of sucrose anabolizing SPS enzyme were significantly increased in S4 and S5 stages compared to the buds, and thereafter a decline was observed in the senescent flowers (S6). Sucrolytic enzymes such as SUS and cytosolic INV also showed a similar trend; a significant increase in enzyme activities were found S4 and S5 petals, which declined in the S6 stage. However, vacuolar INV activity did not show any no significant decline in S6 flowers (Fig. 6A). Histoenzymological technique was used to assess the cINV activities where S4 stage showed maximum activity which gradually declined thereafter (Fig 6B).

**Fig. 6.**
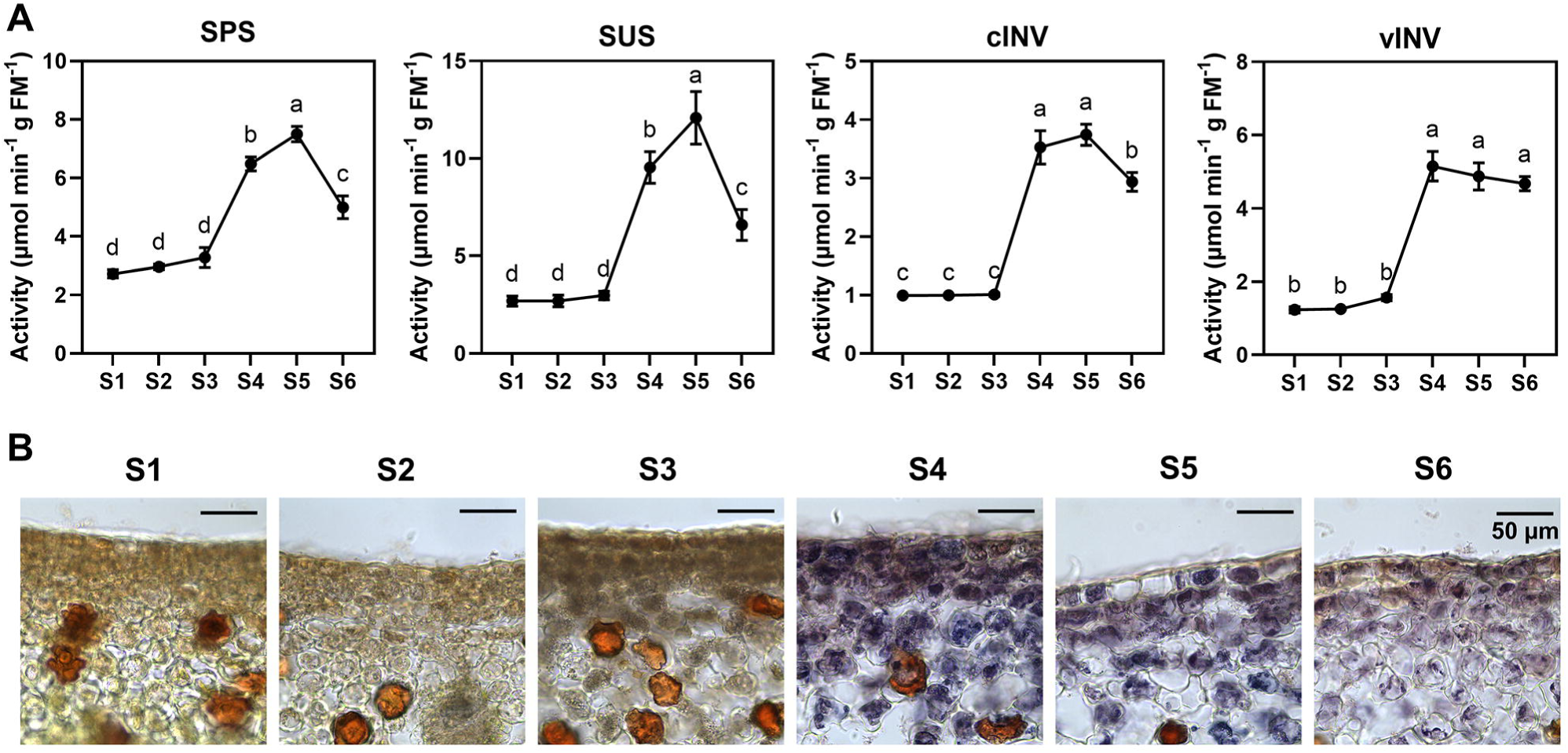
*In vitro* and histoenzymological activities of sucrose metabolizing enzymes. Quantification of *in vitro* activities of SPS, SUS, cINV and vINV activities across different floral maturation stages (A). Histoenzymological detection of cwINV activities in different floral maturation stages (B). The cwINV activity was dectected in petal cross-sections by formazan precipitation. The S4-S6 micrographs show intense blue colouration, indicating presence of invertase activity in corresponding maturation stages. Absence of formazan complex in S1-S3 micrographs indicate low acid wall invertase activity in the early maturation stages of *G. carinata* flowers. n ≥ 3 in all experiments. The data points in line graphs represents mean of at least three biological replicates ± SD. Statistical differences were determined by one way ANOVA and are indicated by different letters (*P* ≤ 0.05).

### Perturbation of petal photosynthesis affects floral maturation

In this study we also addressed how perturbation of photosynthesis in early bud stage (S1) may affect the maturation physiology of the flowers. Perturbed photosynthesis in bud stage caused early onset of senescence-associated-features (Fig. S10) and reduced the floral lifespan (data not shown). We attempted to identify the physiological and metabolic alterations occurred before petal unfurling that led to premature senescence upon flower opening.

A significantly low petal biomass was recorded from the perturbed (PT) buds compared with the control, indicating a reduced growth of buds under perturbed condition of photosynthesis (Fig. 7A, B). Additionally, the PT buds showed markedly low starch and fructus quantity compared to the control (Fig. 7C, D). Histochemical analysis with Lugol’s iodine also showed a low starch abundance in the PT petal tissues (Fig. 7E). *In vitro* measurement of sucrose metabolizing enzymes such as SUS and INVs showed higher activities in the PT petals compared with the normal (Fig. 7F-H). On the contrary, a reduced GAPDH activity was detected in PT petals as compared to normal (Fig. 7I). Metabolomic analyses through GC-MS were conducted in both normal and PT conditions, and the concentration of different metabolites are presented in a heatmap (Fig. 7J). The pairwise comparison between these two experimental conditions are represented in the form of a volcano plot. We observed that the concentration of metabolic sugars such as glucose, fructose and sucrose was significantly higher in the PT buds as compared to the normal (Fig. 7K). Multivariate analysis is presented in form of a scores plot, where a clear separation of the two clusters shows metabolic uniqueness of the two experimental conditions (Fig. 7L).

**Fig. 7.**
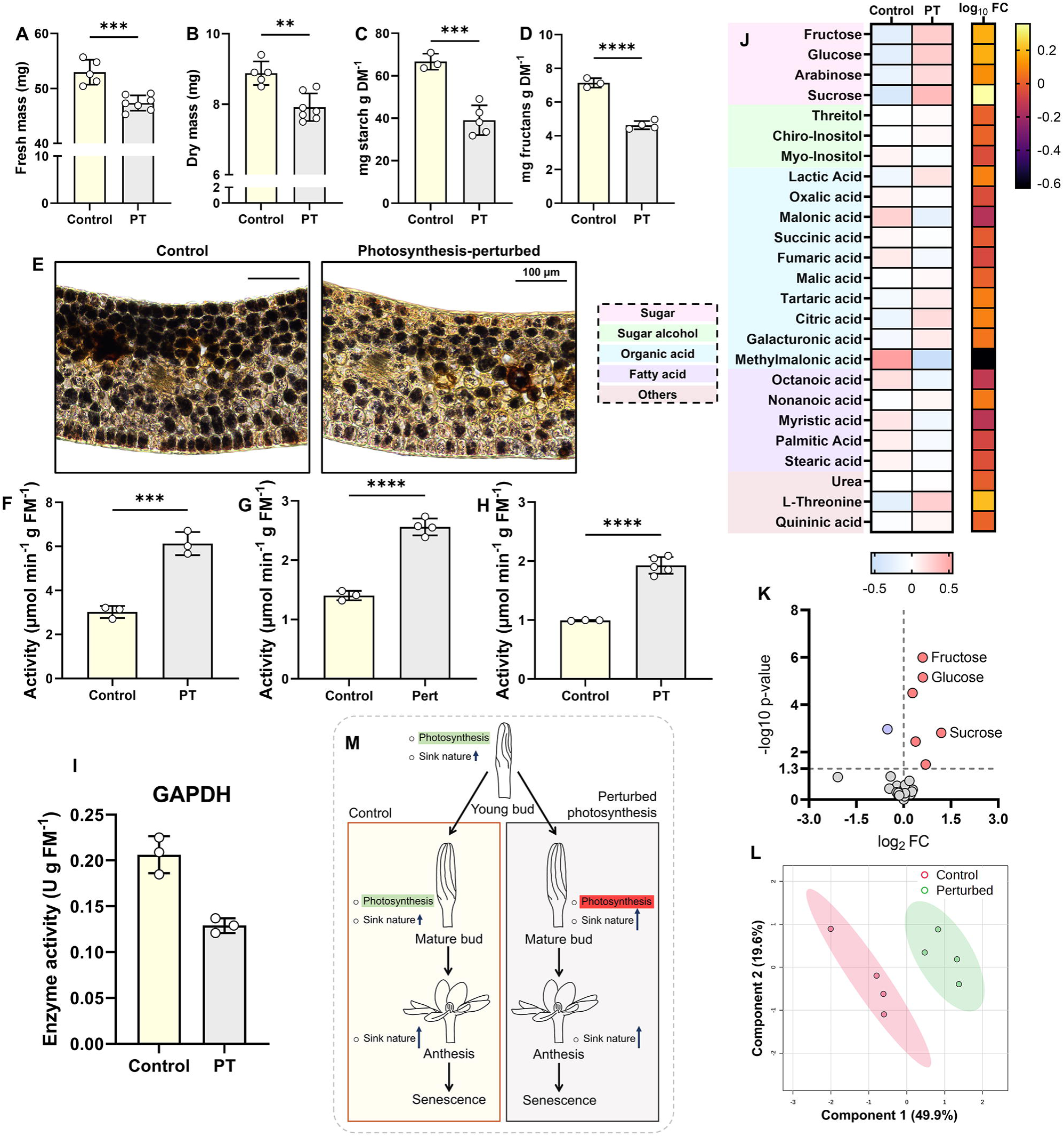
A study showing the consequence of petal photosynthesis perturbation on floral maturation. Fresh and dry masses of flowers (A, B). Quantities of strach and fructans in the flowers (C, D). Histochemical analysis in cryo-sectioned petals with Lugol’s iodine show changes in starch distribution patterns in petals (E). Estimation of *in vitro* activities of SUS, vINV and cINV enzymes (F-H). Estimation of GAPDH enzyme activity (I). Heatmaps showing compositional variation of a few metabolites in control and photosynthesis-perturbed (PT) flowers. Fold change values of the represented metabolites were log_10_-transformed and represented in the associated heatmap (J). Pairwise comparison of metabolite concentrations of the two experimental conditions is represented in volcano plot (K). The red and blue dots show significantly upregulated and downregulated metabolites, respectively, in the PT petals. PLS-DA scores plot showed a clear separation along first two components, indicating significant difference in metabolite composition between the two experimental conditions (L). The first two components accounted for representing 69.5% of the variation. A schematic representation summarizing the major physiological differences among the flowers kept under two different experimental conditions. The flower buds kept for control condition probably has more reliance on petal photosynthesis and show less sink strength compared to PT flowers. *G. carinata* flowers growing under natural condition show elevated sink nature only after flower opening. However, the photosynthesis perturbed flowers increased their sink strength during the bud stages, probably to compensate for the metabolic deficit arising from the absence of photosynthesis (M). n ≥ 3 in all experiments. The data points in line graphs represents mean of at least three biological replicates ± SD. Statistical differences were determined by t-test and are indicated by asterisks (\**P* ≤ 0.05, \*\**P* ≤ 0.01, \*\*\**P* ≤ 0.001 and \*\*\*\**P* ≤ 0.0001).

## Discussion

Flowers in general have fascinated researchers for centuries through their showy displays and fragrant nature. Predictively, the subject of several studies over the past few decades has been to comprehend the specialized metabolic routes that produce the visual and olfactory cues in the flowers. Subsequently, many studies on other aspects of floral physiology surfaced, such as morpho-biochemical, redox changes, and dynamic alteration of various central metabolite classes that occur during floral maturation (Rogers, 2012; Jhanji *et al*., 2025). A recent study also described the metabolomic changes associated with flower anthesis in *Arabidopsis* (Borghi *et al*., 2022). However, our understanding on carbon metabolism that sustains floral physiology and sequentially drives the maturation events during the short floral lifespan, remains poorly understood. Here, we presented a holistic outline of primary metabolism that includes measurement of photosynthetic features and a multilevel analysis of primary metabolite dynamics to find the role of petal photosynthesis and carbon metabolism in floral maturation.

### Photosynthesis is active in *G. carinata* floral buds

Leaves are apparently the photosynthetic organs and other plant parts including reproductive organs, are predominantly function as sink, which depends on the photoassimilates produced by leaves to sustain their growth and physiological processes (Van Camp, 2005; White *et al*., 2016). Nevertheless, several reports have shown that various reproductive structures such as sepals, petals and awns can substantially contribute in assimilating carbon via photosynthesis and supplement the energy demand of reproductive organs (Ashcan and Pfanz, 2003; Leonardos *et al*., 2014; Raven and Griffiths, 2015, Song *et al*., 2024). In *Gagea lutea*, it was reported that excision of floral bracts drastically reduced seed production, while excision of leaves did not show any marked effect on seed development, indicating the role of reproductive whorls in photoassimilation (Sunmonu *et al*., 2013). In our study, we observed the bud stages (S1 to S3) of *G. carinata* flowers were characterized by the presence of well-developed chloroplasts and high quantities of chlorophylls, which represent the main photosynthetically active stages. A reduction in above mentioned photosynthetic features after flower opening indicates loss of photosynthetic abilities in the subsequent maturation stages (S4 to S6). Similar events were reported by Müller *et al*. (2010) in tobacco flowers that showed early carbon assimilating stages, characterized by high quantities of chlorophyll and higher abundance of Rubisco. In *Arabidopsis* flowers, the decline in chlorophyll quantity was associated with the reduction in photosynthesis as the flowers senesced (Pyke and Page, 1998).

Chlorophyll florescence measurements in *G. carinata* flowers also showed consistent results, where the bud stages showed highest photosynthetic activities and the latter maturation stages did not show symptoms of active photosynthesis. A recent study on *Gardenia jasminoides* reported a similar transition, where the floral buds were featured with active photosynthesis while the later stages displayed predominantly heterotrophic features (Ghosh *et al*., 2026). Higher stomatal conductance observed in these photosynthetically active flowe0rs could support efficient CO2 exchange for photoassimilation (Zhang *et al*., 2022). In addition, the higher GAPDH activity in S1 to S3 stages could immediately channelize the photoassimilates to meet the energy and metabolite demands of *G. carinata* floral buds. These findings suggest that the *G. carinata* floral buds are capable of carbon fixation and thus have less reliance on the assimilates produced by the mother plant, which ensure a reduction in reproductive cost. The mother plants thus can support a large number of flowers to ensure reproductive success though efficient carbon allocation (Dorken *et al*., 2025).

### Bud stages are characterized by active growth and accumulation of storage carbohydrates

The floral buds of *G. carinata* are closed green photosynthetic structures which gradually increase their biomass and transition to unfurled flowers upon maturation. Metabolomic studies revealed that these stages (S1 to S3) are characterized by their high sucrose-to-hexose ratio, which gradually declines as the buds mature (Fig. S11). Such ratio had been found to act as metabolic signal that upregulate storage metabolism in plants (Weber *et al*., 1997). Consistent with this model, we also observed in *G. carinata* a gradual accumulation of starch and fructans during the maturation phases (S1 to S3) of floral buds. It is apparent that flowers reserve a considerable amount of storage carbohydrates during bud maturation to meet the physiological and energy demands during flower opening (van Doorn and van Meeteren, 2003; Önder *et al*., 2023; Paul *et al*., 2026).

The metabolic features of the bud stages (S1 to S3) were further substantiated by assessing the activities of a few selected carbohydrate metabolizing enzymes. Previous studies demonstrated starch biosynthesis and biomass accumulation in sink organs are characterized by high activities of invertase(s) and sucrose synthase, the marker enzymes that determine sink-strength (Tang *et al*., 1999; Koch, 2004, Jammer *et al*., 2020). In contrast, our study in *G. carinata* showed low activities of invertase(s) and sucrose synthase from S1 to S3 stages. These findings suggest that *G. carinata* flowers probably rely on locally assimilated carbon for storage and biomass accumulation during these maturation stages. In addition, starch and cell wall biosynthesis enzymes showed higher activities in the growing buds. Subsequently, metabolic precursors of these pathways such as G1P and G6P had higher titer from S1 to S3 stages. Elevated activities of these enzymes and higher concentration of metabolic intermediates explain the gradual increase in starch quantity and biomass as the buds mature. In addition to the hexose sugar phosphorylating activities, the HXK enzymes play a pivotal role in organ development by modulating hormonal interactions and sugar signalling (Granot *et al*., 2013, Granot *et al*., 2014; Sheen, 2014). A study in the past showed fine tuning in photosynthesis metabolism through HXK-dependent glucose sensing in various tissues thus integrate photosynthesis with available carbon status (Garnot *et al*., 2014). Therefore, high activities of HXK in the *G. carinata* buds probably coordinates glucose homeostasis with photosynthetic activities while fulfilling the carbon demand for active growth and storage reserve. Further, through sugar signalling, HXK was shown to prevent the accumulation of miR156, which represses maturation of a plant organs by impeding developmental progression (Garnot *et al*., 2014). No such studies have yet been conducted in flowers that show any possible link between miR156 and HXK during floral maturation, this warrants a mechanistic investigation in near future.

### Flower opening is characterized by energy-extensive metabolism

Comparative metabolomic studies on *G. carinata* petals showed a marked increase in metabolic sugar content (viz. sucrose, glucose and fructose) in anthesis petals (S4) compared to the previous maturation stages. Our observation is consistent with a previous study on Jasminum sambac, where an increased concentration of sugars were reported in the blooming flowers (Ghissing *et al*., 2022). In tobacco flowers, the increased sugar concentration during flower opening was attributed to petal unfurling by regulating the osmotic potential and turgor of petals cells (Stitz *et al*., 2014). Consistent with this concept, increased relative water content was recorded in S4 stage petals. This could be a consequence of increased sugar quantity, which generates turgor subsequently to drive petal opening (Beauzamy *et al*., 2014).

In *G. carinata* flowers, histochemical and in vitro assays revealed a marked reduction in starch, fructans and other soluble carbohydrate quantities during the transition from S4 to S6 stages of maturation (Fig.). Previous studies on floral metabolism also reported a similar trend where the flower opening was characterized by reduction in carbohydrate contents (Waithaka *et al*., 2001; Yap *et al*. 2008; Önder *et al*., 2023a). In plant tissues, α-amylase breakdowns starch and releases simpler sugars (Stanley *et al*., 2005). Increased activity of amylase during *G. carinata* flower opening could be attributed to the reduction in starch and concomitant increase in sugar quantities in anthesis and post-anthesis stages (S4 to S6). Previous studies demonstrated amylase enzymes as the integral part of flower unfurling machinery and thus inhibition of amylase activities were shown to retard flower opening (Rao and Mohan Ram, 1982; Hammond, 1982). A concomitant increase in SUS and INV activities in the S4 stage petals cleaved sucrose into glucose and fructose, thereby accounting to their increased abundance in the petal tissues (Millard *et al*., 2015; Stein and Granot, 2019; Pleyerová *et al*., 2022).

Apart from their function in petal unfurling, sucrose, glucose and fructose are metabolized to provide the energy and metabolic precursors for running specialized metabolism (Dudareva *et al*., 2013; Kutty *et al*., 2021). Therefore, it can be suggested that the higher concentrations of the above-mentioned metabolic sugars probably serve the energy and metabolite demands for volatiles synthesis and its emission in the opened flowers (S4). Floral metabolome analysis revealed an increased trehalose concentration in S4 stage compared with S3 buds in *G. carinata*. Higher trehalose concentration probably fine-tunes the hexose catabolism via a trehalose-6- phosphate mediated signalling to optimally maintain hexose concentrations in these flowers (Zhang *et al*., 2009). Further, *G. carinata* anthesis petals also showed elevated concentrations of energy-rich TCA intermediates such as 2-ketoglutaric acid, succinic acid along with an increased SDH activity. Previous reports on tobacco and Arabidopsis flowers reported similar metabolic reprogramming, where TCA cycle intermediates showed association with floral opening (Stitz *et al*., 2014; Borghi *et al*., 2022). The higher abundance of high-energy TCA intermediates together with increased activity of SDH enzymes in *G. carinata* anthesis petals suggest that an energy demanding metabolism is in operation upon petal unfurling (Busi *et al*., 2011). This hypothesis is further supported by substantial increase in G6PDH enzyme activity at S4 stage. This G6PDH could contribute to the increased energy demand through generation of NADPH during floral maturation (Müller *et al*., 2010). Ultrastructural studies of *G. carinata* S4 petals revealed increased abundance of mitochondria, juxtaposition of ER and mitochondria and presence of prominent Golgi bodies, indicating high metabolic activity in the petal cells (Maiti and Mitra, 2017; Paul *et al*., 2024; Ghosh *et al*., 2026). Taken together, we suggest that the flowers undergo an extensive metabolic reprogramming during flower opening, where they transition from an autotrophic metabolism to an energy-extensive heterotrophic metabolism.

### Changes in metabolic physiology during *G. carinata* floral senescence

At the end of floral lifespan, senescence is triggered by pollination or developmental cues (Dar *et al*., 2014; Borghi and Fernie, 2020). Floral senescence is characterized by marked alterations in the physiochemical traits of cellular membranes, including progressive reduction in membrane integrity (Jhanji *et al*., 2023). In *G. carinata* flowers, a gradual increase in electrolyte leakage and reduced cell integrity could be caused by peroxidation which results in a subsequent loss of cellular homeostasis culminating to cell death (Rogers, 2012; Rogers and Munné-Bosch, 2016; Jhanji *et al*., 2023; Haq *et al*., 2024). TEM micrographs also revealed similar observations, where disintegrating cell membranes were observed in the S6 stage floral petals, substantiating our previous observation on electrolyte leakage and cell death (Ghosh *et al*., 2026).

TEM micrographs showed buckled and thinned cell walls in senescing petal tissues of *G. carinata*. A previous study on carnation flowers revealed a similar observation that reported post-anthesis petals were characterized by their leaning and buckling cell wall (Smith *et al*., 1992). Progressive degradation of cell wall is considered as a character of floral senescence that results into turgor loss and wilting of petals (Shibuya *et al*., 2016). In post-anthesis stages of Rosa flowers, increased activities of cell wall degrading enzymes cause hydrolysis of structural components of cell wall (Önder *et al*., 2023b). In *G. carinata* petals, progressive increase of cellulase and glucanase activities explain the reduction in cellulose quantity in the post-anthesis stages. The increase concentrations of xylitol in S5 and S6 stages could be explained by elevated activities of xylanase in those maturation stages (Brummell *et al*., 2004). Similarly, high concentration of galacturonic acid in post-anthesis stages of G. carinata maturation could indicate substantial degradation of pectin during floral senescence (Brummell *et al*., 2004; Önder *et al*., 2023b). The degradation of cell wall polymers during senescence probably enables the plants to salvage and remobilize the unutilized carbon pool to the newly developing organs (Barnes and Anderson, 2018).

A significant reduction in the concentrations of sucrose, glucose and fructose were recorded in S6 stage petals of *G. carinata*. Previous studies on *Jasminum sambac* and *Clerodendrum chinense* also reported similar observations, where the senescing petals exhibited marked reduction in the quantities of metabolic sugars (Ghissing *et al*., 2022; Paul *et al*., 2026). Reduced metabolic sugar quantity in senescing petals could be a consequence of their extensive utilization in the preceding maturation stages to drive flower opening, scent emission and pigment accumulation in *G. carinata* flowers. Additionally, higher levels of energy-rich TCA intermediates together with increased SDH enzyme activity in S6 petals indicate the flowers probably rely on TCA cycle for the terminal developmental process (Borghi *et al*., 2022). Meanwhile, amino acid profiling of petal tissues and apoplast revealed high abundance of various amino acids in the senescing petals (S6). Senescence associated protein degradation releases the constituting amino acids, which utilized as energy source or can be remobilized to mother plant (Araújo *et al*., 2011; Borghi and Fernie, 2020). To reduce energy cost of the mother plant, unutilized metabolites are usually remobilized to other plant parts (Bieleski, 1995; Borghi and Fernie, 2020). Similarly, reduced activities of sink-determining SUS and INV enzymes in senescent *G. carinata* petals suggest plausible reduction of sink strength to remobilize surplus metabolites (Borghi and Fernie, 2020).

### Role of sucrose metabolism in floral maturation

In most plants, sucrose serves as the primary photoassimilate that is translocated from the source (e.g. photosynthetic leaves) to various sink (e.g. flowers, seeds, roots) tissues through phloem. After sucrose is unloaded into the sink, it is broken down by various sucrolytic enzymes to release hexose sugars, which provide energy for physiological processes and precursors for various metabolic routes (Ruan, 2012). In *G. carinata*, we observed a lower concentration of sucrose in bud stages, which subsequently uplifted in S4 stage, and thereafter declined in senescing flowers. In vitro assay of a sucrose biosynthetic enzyme SPS showed corresponding trend with sucrose concentration, where notable increase in enzyme activity was recorded in S4 stage petals as compared to the previous stages. However, a significant decline of SPS activity was observed in the terminal maturation stage (S6). It was further reported that SPS work tandemly with sucrose efflux transporters in Arabidopsis to maintain the sucrose titer in plant organs (Chen *et al*., 2012). These observations suggest that through SPS activities the petal tissues possibly maintain the necessary sucrose pool required during the different stages of floral maturation. However, the role of this enzyme in sink determination remained elusive and warrants further investigations (Micallef *et al*., 1995, Ruan, 2014). In contrast, SUS, a sucrose- breakdown enzyme is widely considered as the biochemical hallmark of sink tissues (Gessler, 2021). In sink tissues such as fruits and seeds, SUS activity ensures optimal growth and accumulation of storage carbohydrates. In G. carinata, a significant increase in SUS activity was found upon flower opening after losing their photosynthetic abilities. This suggests that the flowers need a strong sink demand as they entered into their energy demanding stages. Thus, elevated SUS activities from S4 to S6 stages enable the petal tissues to establish an efficient supply of sucrose in opened flowers.

Similar to SUS, invertases (INVs) represent another type of sucrolytic enzymes that ensure a negative concentration gradient of sucrose to maintain the sink strength in non-photosynthetic organs. Based on the subcellular localization and pH optima, the INVs are classified into three groups – cell wall (apoplasmic), cytosolic and vacuolar INVs (Sturm, 1999). The cell wall INVs help to breakdown sucrose molecules in the apoplast and maintain a low apoplasmic sucrose gradient, which favours continuous phloem unloading in the sink tissues (Sherson *et al*., 2003; Yan *et al*., 2019). In *G. carinata* flowers, the cwINV activities increased significantly upon flower opening (S4) as compared to early maturation stages (S1 to S3); a gradual decline of cwINV activities was observed until S6 stage. Concurrently, we found a similar trend in apoplastic sucrose, glucose and fructose concentration, indicating sucrose breakdown in floral apoplast. Similar observation was reported in young tomato fruits, where a fruit specific cwINV facilitates sucrose breakdown for sustained phloem unloading and maintain sink strength (Jin *et al*., 2009). The activities of cINV and vINV enzymes also showed a similar trend as observed with cwINV in *G. carinata* flowers. These observations suggest a probable synergistic role of these enzymes to set a sink demand in the floral tissues. Additionally, vINV breaks down sucrose into fructose and glucose and doubles the osmotic demand of cell vacuoles (Ruan *et al*., 2010). Such changes in osmotic potential drives cell expansion, and thus higher vINV activities could be correlated with increased relative water content and petal unfurling in S4 stage of *G. carinata* (Beauzamy *et al*., 2014). In conclusion, the survey on sucrose metabolism in G. carinata flowers showed that the sink-markers prominently increase upon flower unfurling. Therefore, upon considering the photosynthetic abilities and central metabolite status throughout the floral lifespan, we conceived that the *G. carinata* flowers transitioned from autotrophic to heterotrophic metabolism during their maturation.

### Bud photosynthesis is essential for floral maturation

A comparison between normally developed and PT buds revealed significant alterations in their physiological and metabolomic features. The marked reduction in biomass and storage reserves in PT buds suggests that bud photosynthesis plays a critical role during the active growth phase. During their early growth, floral buds rapidly increase their biomass and storage reserve necessary for subsequent maturation processes. Several studies demonstrated that photosynthesis in reproductive parts is correlated with carbon accumulation in the sinks (Wingler and Soualiou, 2025). It can be hypothesized that limiting photosynthesis in floral buds can reduce the accumulation of storage compounds that lead to lower carbon currency available to support floral physiology during the late maturation stages. Consequently, flowers unfurled from the PT buds showed shortened lifespan. A similar phenomenon was reported in tobacco flowers where photosynthesis-perturbed flowers showed early symptoms of senescence (Müller *et al*., 2010). In contrast to our observation with *G. carinata* flowers, under similar experimental conditions, the young *G. jasminoides* PT buds did show any growth afterwards (data not shown).

Comparative metabolomic analysis between control and PT buds revealed a significant upregulation of metabolic sugars such as glucose, fructose, and sucrose in the PT buds. The upregulation of these sugars could be attributed to the higher activities of sucrose-metabolizing enzymes (such as SUS and INVs), which cleaved sucrose into its constituting hexoses viz. glucose and fructose. Concomitantly, higher activities of SUS and INV enzymes indicate that bud tissues prematurely acquire their sink nature following perturbation of photosynthesis. Alongside, the elevated accumulation of hexoses lowers the sucrose-to-hexose ratio, causing reduced biomass and storage accumulation in PT buds (Weber *et al*., 1997; Ruan, 2014). Furthermore, the elevated glucose titer in the PT buds can be sensed by HXK enzymes, which are known to impede photosynthesis in presence of glucose (Garnot *et al*., 2014). Collectively, our findings suggest that the *G. carinata* floral buds act as photosynthetically active source tissues, characterized by low sink determining enzyme activities, low glucose levels and a high sucrose-to-hexose ratio under normal maturation conditions. Perturbation of bud photosynthesis impairs the aforementioned metabolic balance and prematurely establishes sink metabolism at the expense of bud growth, which demonstrate that bud photosynthesis is essential for maintenance of normal maturation physiology of flowers.

## Conclusion

This study provides a comprehensive insight into the dynamics of carbon metabolism in *G. carinata* flowers and how it underpins the physiology of flower maturation. We found that the floral buds are photosynthetically active and petal photosynthesis significantly contributes in accumulating carbon reserve, essential for later stages of floral maturation. Furthermore, while metabolic events are considered to be orchestrated by floral maturation events, our study suggests floral metabolic frame may also influence the maturation events. Flowers of *G. carinata* after opening, showed an energy-extensive metabolism that probably ensures the flowers to unfurl and support energy and metabolite requirements of scent and pigment metabolism. As the flowers move towards senescence, activities of several cell wall degrading enzymes increased. This suggests that the carbon stored in cell wall is released again in the cytosol, which may support metabolism or can be remobilized to other plant parts. Lastly, our study empirically showed that autotrophic buds possess low sink strength, and upon unfurling they enter into a heterotrophic metabolism and converted into strong sinks. Further reduction in INVs and SUS enzyme activities lower the sink strength probably to salvage the unutilized carbon pool through remobilization (Fig. 8).

**Fig. 8.**
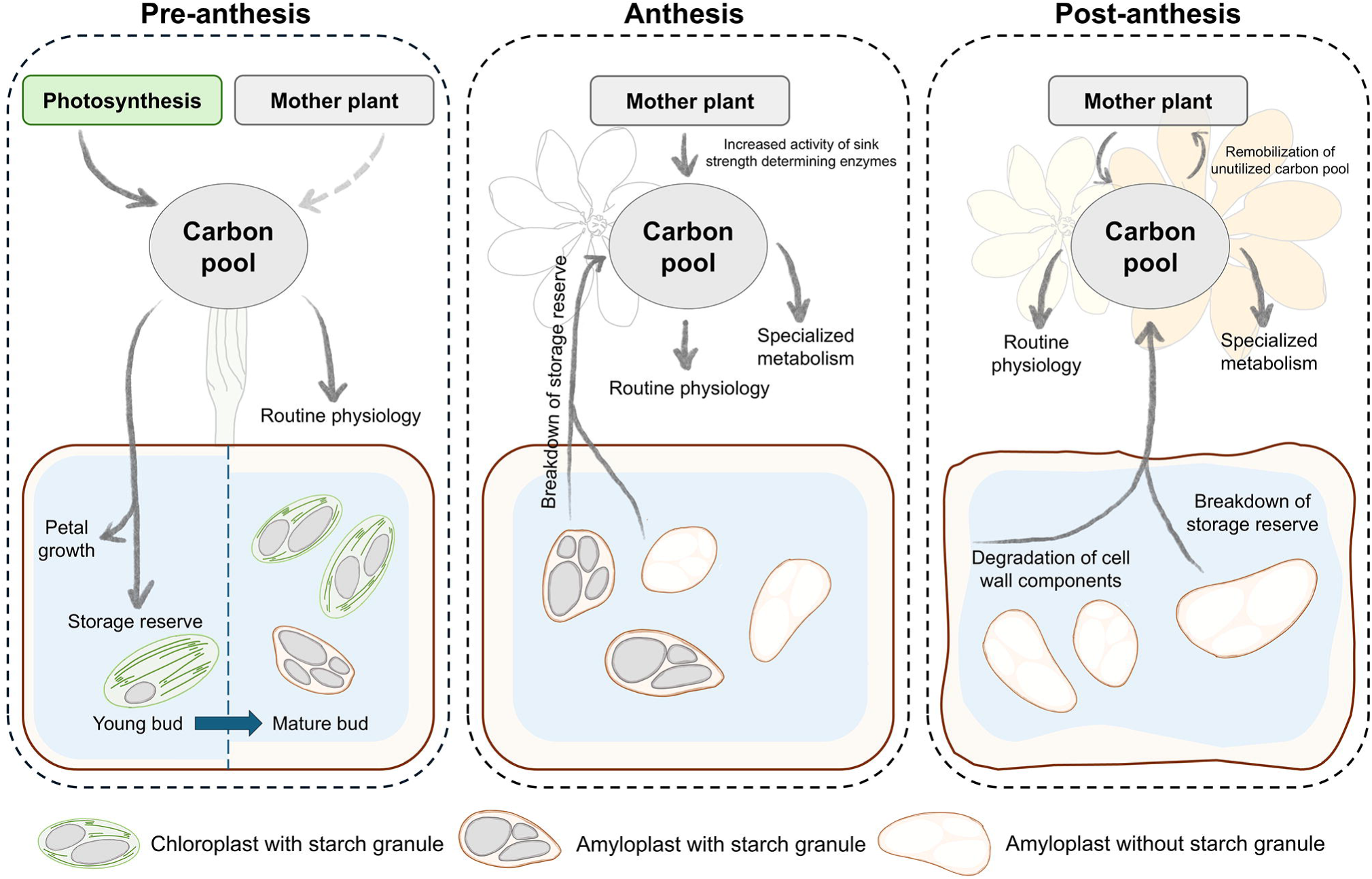
A scheme depicting major metabolic events that occur during floral maturation. The floral buds show photosynthetic abilities and low sink strength, indicating the occurance of autotrophic metabolism in *G. carinata* floral buds. During maturation, the floral buds show active growth and accumulate substantial amount of storage reserve. During anthesis, significant amount of storage reserve gets utilized, probably to propel flower opening and high energy demand associated with scent volatile synthesis and emission. Loss of photosynthetic abilities and increased sink-strength of anthesis flowers signifies the flowers probably transition from autotrophic to heterotrophic mode of metabolism during flower opening. As the flowers reach senescence stage, storage reserves get significantly depleted and cell wall degradation also start releasing monomeric sugars, which contributes to terminal metabolic events. A reduction of sink strength in senescence stage could enable flowers to remobilize their unutilized carbon pool back to the mother plant.

## Supplementary data

### Supplementary method S1. GC-MS parameters

**Supplementary Fig. S1.** Petal stomatal conductance

**Supplementary Fig. S2.** SEM micrographs of petal

**Supplementary Fig. S3.** TEM micrographs of petal

**Supplementary Fig. S4.** Dendrogram showing hierarchical clustering of metabolome data

**Supplementary Fig. S5.** Quantification of G1P and G6P

**Supplementary Fig. S6.** *In vitro* activities of G6PDH, GAPDH and SS

**Supplementary Fig. S7.** Histoenzymological detection of isocitrate dehydrogenase activity

**Supplementary Fig. S8.** Histoenzymological detection of succinate dehydrogenase activity

**Supplementary Fig. S9.** PAS staining of petal sections

**Supplementary Fig. S10.** Electrolyte leakage and cell integrity measurements of PT petals

**Supplementary Fig. S11.** Sucrose-to-hexose ratio in different maturation stages

## Acknowledgement

The authors acknowledge the technical personnel of Sophisticated Analytical Instrumentation Facility, All India Institute of Medical Sciences, New Delhi for providing kind support in TEM sample processing and image acquisition.

## Author contribution

RG: Conceptualization, methodology, investigation, formal analysis, data curation, validation, writing – preparation of original draft; AM: conceptualization, funding acquisition, resources, supervision, writing – revisions and finalization of manuscript.

## Data availability

All data related to this article have been presented in the main manuscript and the supplementary file. Additional information will be provided upon reasonable request.

